# The effect of resource limitation on the temperature-dependence of mosquito population fitness

**DOI:** 10.1101/2020.05.29.123315

**Authors:** Paul J. Huxley, Kris A. Murray, Samraat Pawar, Lauren J. Cator

## Abstract

Laboratory-derived temperature dependencies of life history traits are increasingly being used to make mechanistic predictions for how climatic warming will affect vector-borne disease dynamics, partially by affecting abundance dynamics of the vector population. These temperature-trait relationships are typically estimated from populations reared on optimal resource supply, even though natural populations of vectors are expected to experience variation in resource supply, including intermittent resource limitation. Using laboratory experiments on the mosquito *Aedes aegypti*, a principal arbovirus vector, combined with stage-structured population modelling, we show that low-resource supply significantly depresses the vector’s maximal population growth rate across the entire temperature range (22-32°C) and causes it to peak at a lower temperature than at high-resource supply. This effect is primarily driven by an increase in juvenile mortality and development time, combined with an exaggerated decrease in adult size with temperature at low-resource supply. Our study suggests that projections of vector abundance and disease transmission based on laboratory studies are likely to substantially underestimate how resource supply can modulate the temperature-dependency of population-level fitness through its influence on juvenile survival and development time. Our results provide compelling evidence for future studies to consider resource supply when predicting the effects of climate and habitat change on disease vectors and transmission.

## INTRODUCTION

The global burden of human, animal, and plant vector-borne diseases has increased substantially in recent decades [1,2]. The transmission patterns of these diseases are strongly linked to spatio-temporal distribution and abundance of their vectors [3,4]. Therefore, there is growing concern that climate and land-use change coupled with rapid globalization may shift the distributions and abundances of vector species and thus, the diseases they transmit [5,6]. However, we currently lack a mechanistic understanding of how changes in multiple abiotic environmental drivers affect the abundance of disease vectors [7-9].

Because most disease vectors are small ectotherms, interest is growing rapidly in the effects of environmental temperature in particular on their population fitness [5,6,10–15]. Biological rates (“functional traits”, such as metabolic, development, and mortality rate [7]) of ectotherms increase approximately exponentially with temperature up to some optimum before declining to zero [16]. Therefore, a number of recent studies have used laboratory data on the thermal responses of functional traits of vectors to predict how temperature will affect vector abundance and disease transmission in the field, leading to interesting new insights [8]. However, laboratory-derived temperature-trait relationships are generally measured in populations reared on optimal resource supply. Yet, natural populations of vectors are expected to experience variation in resource supply, including intermittent resource limitation.

Indeed, along with temperature, resource supply is another ubiquitous environmental driver that is expected to limit the fitness of vector populations in nature [17-19]. Moreover, temperature and resource supply are expected to act interactively [9,20]. The primary reason for this, is that while the energy cost of somatic maintenance, growth and ontogenetic development of individuals generally increases with temperature [21,22], the ability to meet this increasing demand depends on resource supply. If the resources available to an individual do not keep pace with increasing energy requirements, its growth, development, and survival would be compromised. Ultimately, this should negatively affect fitness, with the severity of these effects increasing with temperature. While the importance of resource supply in mediating the effect of temperature on population abundance may seem obvious, this problem remains largely unresolved theoretically and empirically, not just in vector-borne disease research, but in thermal ecology in general. For example, Ecological Metabolic Theories (including the Metabolic Theory of Ecology (MTE) and Dynamic Energy Budget (DEB) theory), which seek to link organismal metabolic rates to ontogenetic and population growth, generally assume that resource supply is not a limiting factor [21-24].

Additionally, resource supply can also interact with temperature to affect population fitness of ectotherms by determining size at maturity. Generally, size at maturity decreases with rising temperature (the size-temperature rule [25], which also applies to disease vectors, such as mosquitoes [26]). However, the size-temperature rule too remains largely untested under resource limitation in disease vectors and other ectotherms [25,27]. For vectors specifically, female size is demographically and epidemiologically important because it is associated with longevity, fecundity, and biting behaviour [28,29].

In general, the question of whether and how temperature and resource supply may together modulate disease transmission through underlying traits remains open. Here we seek to fill this gap in knowledge by investigating the effect of realistic variation in resource supply on the temperature-dependence of population-level fitness in *Aedes aegypti*, a principal mosquito vector of human arboviruses (e.g. dengue, yellow fever and zika; [38]). Because resource competition between larvae is expected to be a major regulator of adult mosquito abundance, many studies have examined how resource supply and larval density interact to affect fitness [30–33], while others have investigated the effect of resource supply and larval density separately [34,35]. However, none of these studies have considered environmental temperature. On the other hand, studies that have considered temperature have not examined how the effects of temperature and resource supply together affect fitness through traits [36,37]. By taking a trait-based approach, we seek to gain general, mechanistic insights into how resource availability and temperature may together affect the abundance of disease vectors in the field.

## METHODS

To investigate the effects of temperature and resource supply on mosquito life history, we employed a 3×2 factorial design comprised of three temperatures (22, 26, and 32°C) and two resource supply levels: 0.1 (low-resource supply) and 1 mg larva^-1^ day^-1^ (high-resource supply). We conducted a preliminary assay to determine the adequacy of this replication level to detect statistically significant effect sizes (electronic supplementary material, Tables S4, S5). These experimental temperatures span the range of average annual temperatures [39] that this strain of *Ae. aegypti* is likely to experience in the wild (F16-19 originating from Fort Meyer, FL; [40]). This low-resource supply level was chosen because previous work has found that it is the highest resource limitation that can be applied to this species without resulting in complete juvenile mortality [17,18]; a level of limitation that might be expected in wild populations. We also determined that the low-resource level was appropriate with a preliminary assay (electronic supplementary material, Tables S4, S5). The high-resource supply level corresponds to the upper mid-range of the high-resource supply levels used in Arrivillaga and Barrera [17] and Barrera et al. [18], and is consistent with the levels of resource supply commonly used in laboratory studies on this species [36,41].

Batches of approximately 300 *Ae. aegypti* eggs were randomly assigned to one of the three experimental temperatures and immersed in plastic tubs containing 150 ml of tap water. Each tub was provided with a pinch of powdered fish food (Cichlid Gold®, Hikari, Kyrin Food Industries Ltd., Japan) to stimulate overnight hatching. The tubs were then submerged in water baths (Grant Instruments: JAB Academy) set at either 22, 26, or 32°C. Water baths were situated in a 20°C climate-controlled insectary with a 12L:12D photoperiod and 30 minutes of gradual transition of light levels to simulate sunrise and sunset. On the following day, first instar larvae were separated into cohorts of 30 and held in tubs containing 150 ml of water. We created three replicate tubs per treatment (90 individuals/treatment). Low-resource supply treatments were provided 3 mg of food and high-resource supply treatments received 30 mg. Thereafter, resource levels were adjusted daily according to the number of living individuals in each tub prior to feeding each day such that resource levels were maintained at an approximately constant level during the juvenile lifespan. Rearing tubs were cleaned and refilled with fresh tap water daily. Water volumes were also adjusted daily in accordance with mortality to maintain larval density (0.2 larvae × ml^-1^). Figure S1 is a schematic of the experimental design and the traits measured.

### Fitness calculation

To calculate population-level fitness, we used our data to parameterise stage-structured matrix projection models [42], which describe change in a population over time:

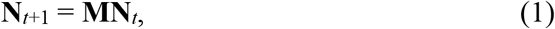

where **N***t* is a vector of abundances in the stage classes at time *t* and **M** is the population projection matrix. The first row of **M** is populated by daily fecundity (the number of female offspring produced per female at age *i*). The sub-diagonal of **M** is populated with the probabilities of survival from age *i* to age *i*+1. Multiplying the transition matrix (**M**; eqn. 1) and stage-structured population size vector (**N***t*; eqn. 1) sequentially across time intervals yields the stage-structured population dynamics. Once the stable stage distribution of the abundance vector is reached, the dominant eigenvalue of the system is the finite population rate of increase (*λ*) [42]. Then, the intrinsic rate of population growth is

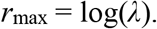

This is a population’s inherent capacity to reproduce, and therefore a measure of population-level fitness [23,43,44]. Negative *r*_max_ values indicate decline and positive ones, growth. The projection matrices were built and analysed using the `popbio` R package [45,46].

### Parameterisation

#### Immature development time and immature and adult survival proportions

Matrix survival elements (the sub-diagonal of the matrix **M**; eqn. 1) were populated with continuous survival proportions estimated using the Kaplan-Meier survival function in the `survival` R package [45,47]. We assumed life stage duration (i.e. larva-to-pupa-to-adult) was the mean duration of transitioning into and out of that stage, and a fixed age of adult emergence at the mean age of emergence. Adult survival elements were populated with the Kaplan-Meier proportions. Hatching-to-adult development times were calculated by recording the day and time that egg eclosion, pupation and adult emergence occurred for each individual. Upon pupation, mosquitoes were held in individual falcon tubes containing 5 ml of tap water. This enabled pupa-to-adult development durations and the lifespans of individual starved adults to be recorded. Starvation forces adults to metabolise the nutritional reserves accumulated during larval development, so starved lifespan should increase with body size. Therefore, starved adult lifespan is a useful indicator of the carry over effects of temperature and resource availability in the larval habitat [48,49].

#### Daily fecundity rate

The use of scaling relationships between fecundity and size is common in predictions of population growth in *Aedes* mosquitoes [50,51]. A detailed description of our method for estimating fecundity is provided in the electronic supplementary material (Fig. S2). Briefly, we measured wing length as a proxy for body size, and estimated lifetime fecundity using previously published datasets on the temperature- and resource supply-dependent scaling between wing length and lifetime fecundity [41,49]. Daily fecundity rate is required for the first row of **M** (eqn. 1), so lifetime fecundity was divided by lifespan and multiplied by 0.5 (assuming a 1:1 male-to-female offspring ratio) to give temperature-specific individual daily fecundity.

### Parameter sensitivity

We used the delta method to approximate 95% confidence intervals (CIs) for our fitness calculations [42,52] to account for how uncertainty in survival and fecundity estimates is propagated through to the *r*_max_ estimate. This method requires the standard errors of the survival and fecundity element estimates. For survival, we used the standard errors estimated by the Kaplan-Meier survival function in the `survival` R package. For fecundity, we calculated the standard errors of the mean daily fecundity rates (electronic supplementary material, Table S2) for each treatment using the `Rmisc` R package [53]. As an additional sensitivity analysis, we recalculated fitness using the upper and lower 95% CIs of the exponents for the scaling of wing length and lifetime fecundity (Fig. 3, electronic supplementary material, Fig. S2).

**Fig. 1.**
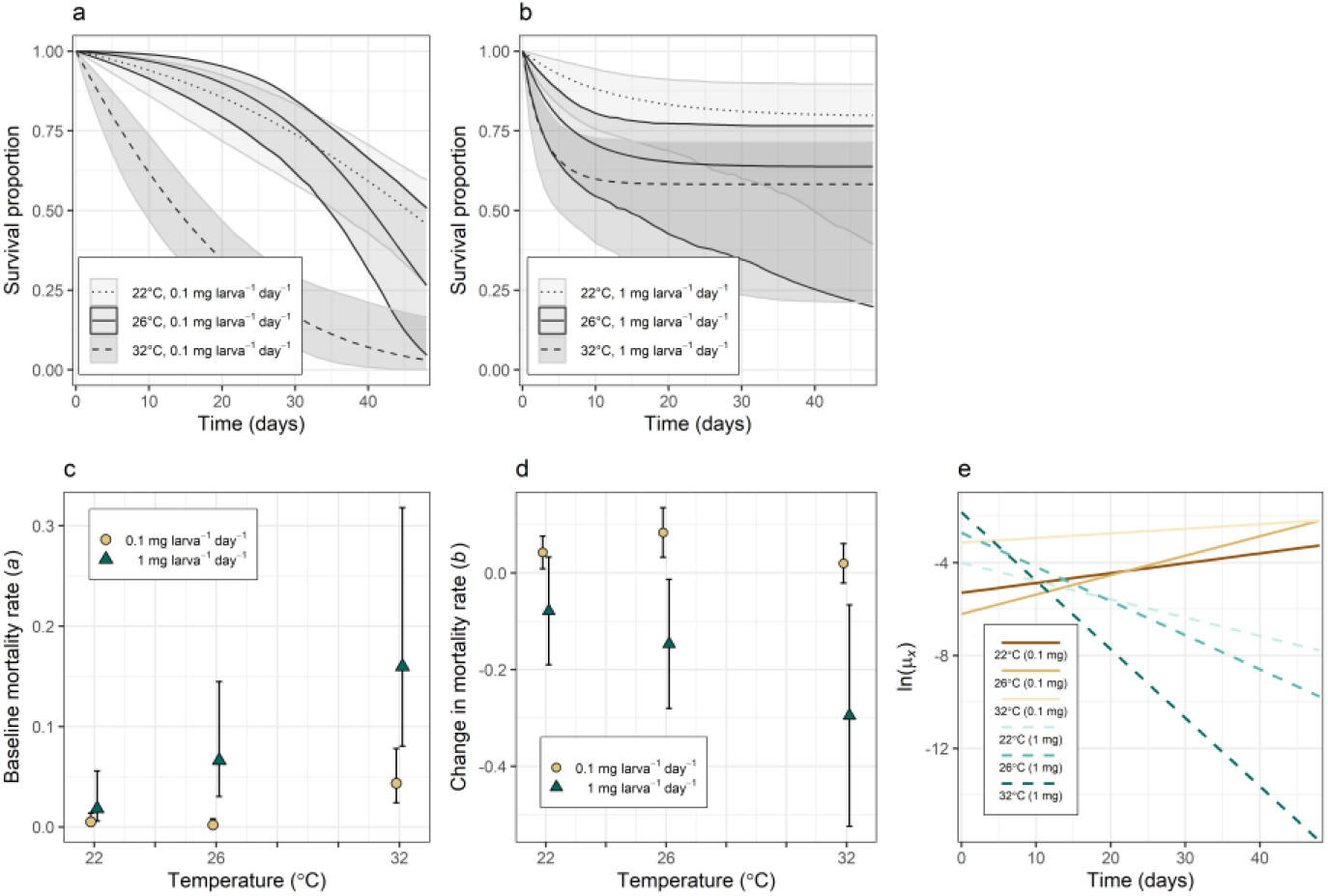
The effect of resource supply on the temperature-dependence of juvenile survival. Survival curves at **(a)** low- and **(b)** high-resource supply by temperature with 95% confidence bounds. **c**, Baseline mortality rates, *a*, by resource supply level across temperatures with 95% CIs. Mortality rates were significantly lower at low-resource supply than at high-resource supply as temperatures increased from 22 to 32°C (95% CIs at 26 and 32°C do not overlap). **d**, Change in mortality rate trajectories, *b*, by resource supply level across temperatures with 95% CIs. Rate trajectories were significantly lower at high-resource supply than at low-resource supply as temperatures increased from 22°C (95% CIs at 26 and 32°C do not overlap). **e**, Logged daily mortality rates (ln(*μ_x_*) = ln(*a*) + *bx*) show how mortality rates at low-resource supply started low and increased with time; at high-resource supply they started high and decreased with time.

**Fig. 2.**
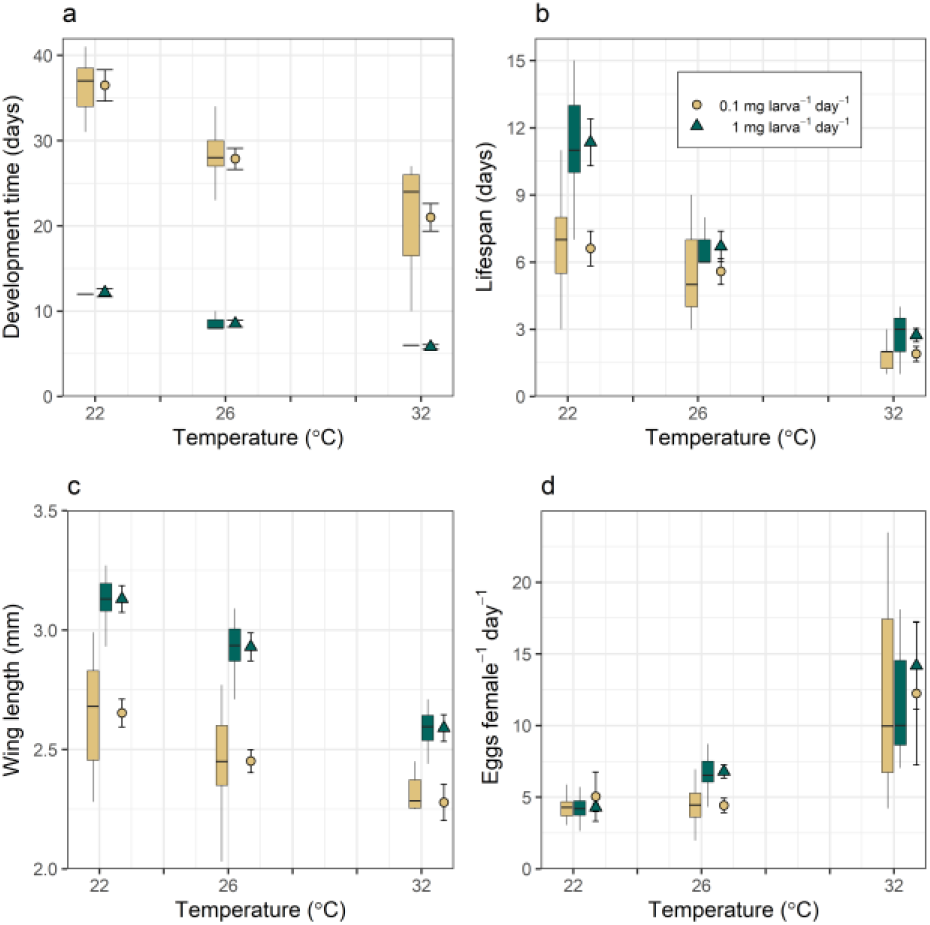
The combined effect of temperature and resource supply level on *Ae. aegypti* life history traits. **a-c**, Resource supply significantly modulated the effect of temperature on all directly measured traits, except adult lifespan at 26°C (**b**). Symbols denote the GLM-estimated means with 95% CIs calculated from the standard errors (electronic supplementary material, Table S2) for the resource supply levels at each temperature (circles, low-resource supply; triangles, high-resource supply). The resulting ANOVAs of the GLMs for each trait are presented in Table S1 (electronic supplementary material). **d**, Predicted fecundity increased significantly with temperature at both resource levels (non-overlapping 95% CIs). The extent to which fecundity increased with temperature and resource level was only significant at 26°C. The numbers of females that survived to adulthood (*n*) in each treatment were: 22°C at low-resource supply *n* = 23, at high-resource supply *n* = 37; 26°C at low-resource supply *n* = 29, at high-resource supply *n* = 30; 32°C at low-resource supply *n* = 10, at high-resource supply *n* = 27. Boxplot horizontal lines represent medians. Lower and upper hinges are the 25th and 75th percentiles. Upper whiskers extend from the hinge to the largest value no further than 1.5 × inter-quartile range (IQR) from the hinge. The lower whisker extends from the hinge to the smallest value at most 1.5 × IQR of the hinge.

**Fig 3.**
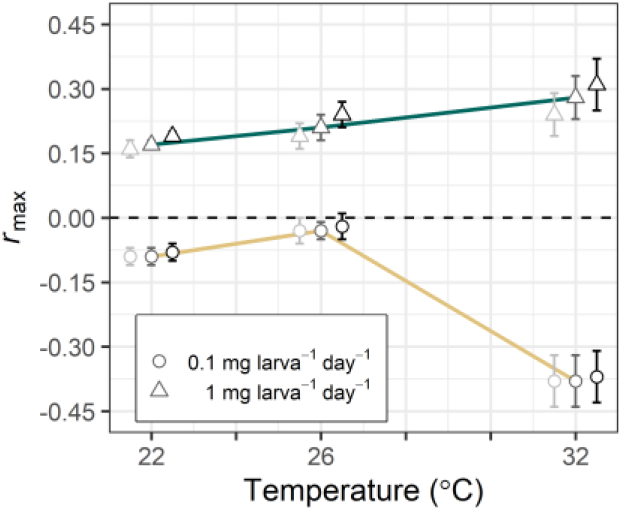
Population-level *Ae. aegypti* fitness (*r*_max_) by resource supply (0.1 (low) or 1 mg larva^-1^ day^-1^ (high)) across temperatures. Fitness estimates for each treatment, with 95% confidence intervals (CIs). The three data points for each treatment represent *r*_max_ estimated using the 95% CI bounds of the exponents for the scaling of lifetime fecundity with wing length (electronic supplementary material, eqn. S1, Fig. S2). The lightest greyscale hue estimates derive from the lower 95% CIs, the midrange hue estimates with trend lines derive from the slopes and the darkest hue derive from the upper 95% CIs.

### Elasticity analysis

Elasticities were used to quantify the proportional contributions of individual life history traits to *r*_max_. Elasticity, *e*_ij_, measures the proportional effect on *λ* of an infinitesimal change in an element of **M** (eqn. 1) with all other elements held constant (the partial derivative) [54,55]. This partial derivative of *λ*, with respect to each element of **M**, is *S_ij_ = ∂λ/∂a_ij_* = *V_i_W_j_* with the dot product 〈**w**, **v**〉 =1. Here, **w** is the dominant right eigenvector (the stage distribution vector of **M**), **v** is the dominant left eigenvector (the reproductive value vector of **M**), and *a_ij_* is the *i*×*j*^th^ element of **M**. Elasticities can then be calculated using the relationship: *e*_ij_ = *a_ij_λ × S_ij_*. Multiplying an elasticity by *λ* gives the absolute contribution of its corresponding *a_ij_* to *λ* [54,55]. Absolute contributions for juvenile and adult elements were summed and changed proportionally to quantify the sensitivity of *r*_max_ to these traits.

### Statistical analyses

All statistical analyses were conducted using R [45]. The trait data (development time, lifespan and wing length) were nonlinear, positive and right skewed, so we used full factorial generalized linear models (GLM) with gamma distributions and identity link functions (predictor effects were considered additive) to determine the significance of each predictor on the thermal response of each of these traits. Replicate was included in these GLMs as a fixed effect.

We investigated the effect of temperature and resource supply on juvenile mortality rate by fitting a set of candidate distributions (exponential, log-logistic, Gompertz and Weibull) to the survival data with R package `flexsurv` [56]. The Gompertz survival function was the best fit to these data according to the Akaike Information Criterion (AIC) (electronic supplementary material, Table S3). The final mortality model was obtained by dropping terms from the full model (consisting of temperature × resource supply + replicate as fixed effect predictors). If removing a term did not improve model fit (ΔAIC > −2), it was removed from the full model (electronic supplementary material, Table S3). For each treatment, maximum likelihood methods executed in `flexsurv` estimated mortality parameters (and their 95% CIs) of the Gompertz model, *μ_x_* = *ae^bx^*. where *a* is the baseline morality rate, and *b* is the change in mortality rate with time. These parameter estimates were then used to determine the significance of the effects of temperature and resource supply on juvenile mortality.

## RESULTS

All trait responses varied significantly with temperature and resource supply, with a significant interaction between the two environmental variables (Figs. 1, 2; electronic supplementary material, Tables S1, S2). Thus, the realised effect of temperature on trait responses was consistently and significantly mediated by resource supply.

At low-resource supply, daily juvenile mortality rates, *μ_x_*, increased with time at all temperatures, whereas, at high-resource-supply, they decreased with time (Fig. 1e). Baseline juvenile mortality rates, *a*, were significantly lower at low-resource supply than at high-resource supply as temperatures increased from 22 to 32°C (non-overlapping 95% CIs at 26 and 32°C; Fig 1c). Mortality rate trajectories, *b*, were significantly lower at high-resource supply than at low-resource supply as temperatures increased from 22 to 32°C (non-overlapping 95% CIs at 26 and 32°C; Fig 1d).

Development time varied significantly with the interaction between temperature and resource supply (ANOVA; *F*_2,0.75_ = 24.11, *p*<0.001; electronic supplementary material, Table S1). Whereas development time decreased both at warmer temperatures and at high-resource supply, the decrease with temperature was greater at low-resource supply than at high-resource supply. At low-resource supply, development time decreased by 15.45 days as temperatures increased from 22 to 32°C, whereas at high-resource supply, it decreased by 6.38 days across this range (Fig. 2a, electronic supplementary material, Table S2).

Adult lifespan varied significantly with the interaction between temperature and resource supply (ANOVA; *F*_2, 2.41_ = 14.95, *p*<0.001; electronic supplementary material, Table S1). Although lifespan decreased both at warmer temperatures and at low-resource supply, the decrease with temperature was greater at high-resource supply than at low-resource supply. High-resource supply lifespan decreased by 8.89 days, whereas low-resource supply lifespan decreased by 4.71 days as temperatures increased from 22 to 32°C (Fig 2b, electronic supplementary material, Table S2).

The interaction between temperature and resource supply resulted in significant variation in size at maturity (wing length) between resource levels (ANOVA; *F*_2,0.03_ = 4.36, *p*=0.01; electronic supplementary material, Table S1). Adult size decreased both at warmer temperatures and at low-resource supply, though the decrease with temperature was greater at high-resource supply than at low-resource supply. At low-resource supply, size decreased by 0.37 mm as temperatures increased from 22 to 32°C, while at high-resource supply, size decreased by 0.54 mm (Fig 2c, electronic supplementary material, Table S2).

### Population fitness (*r*_max_)

Resource limitation depressed *r*_max_ to negative values at all temperatures, with a unimodal relationship of *r*_max_ with temperature (Fig. 3, electronic supplementary material, Table S2). Low-resource supply *r*_max_ increased from −0.09 at 22°C to −0.03 at 26°C and then decreased acutely to −0.38 at 32°C. In contrast, at high-resource supply, *r*_max_ was always positive and increased monotonically with temperature from 0.17 at 22°C to maximal growth (0.28) at 32°C.

### Elasticity analysis

Juvenile survival was the most important contributor to *r*_max_ (electronic supplementary material, Fig. S3). For example, at low-resource supply at 32°C, a 0.5 proportional increase in juvenile survival would almost halve the rate of decline from −0.384 to −0.204 (electronic supplementary material, Fig. S3a). In contrast, for the same treatment, a proportional increase of the same magnitude for adult survival would increase *r*_max_ from −0.384 to −0.379 (Fig. S3b), and fecundity would increase *r*_max_ from −0.384 to −0.376 (Fig. S3c). This underlines how the temperature-dependence of *r*_max_ derives mainly from how resource supply level impacts juvenile mortality and development, which determine the number of reproducing individuals and the timing of reproduction, respectively. Fecundity and adult survival, on the other hand, have relatively negligible effects on *r*_max_, which suggests that the carry over effect of reduced size at maturity on *r*_max_ is relatively weak.

## DISCUSSION

Our results show that juvenile resource regimes can have far-reaching effects on the temperature-dependence of population-level fitness, *r*_max_. Differences between the thermal response of traits at low-versus high-resource supply resulted in a marked divergence of the temperature-dependence of *r*_max_ between the two resource levels (Fig. 3). At low-resource supply, fitness was negative throughout and unimodal (declining steeply from 26 to 32°C). This indicates that population fitness becomes increasingly, and non-linearly constrained by resource limitation as temperatures increase. Fitness at high-resource supply was positive and monotonically increasing. This lack of unimodality indicates that the temperature at which fitness peaks is higher than 32°C when resources are abundant.

The elasticity analysis shows that the primary mechanism underlying the divergent temperature-dependence of *r*_max_ across resource levels is decreased juvenile survival at low resources, which decreased population-level reproductive output (Figs. 1, 3). At low-resource supply, the daily mortality rate started low and then increased over time, while at high-resource supply, it started high and then decreased to very low levels (Fig. 1e). The effect of resource limitation on fitness was further compounded by the increase in juvenile development time (Fig. 1a), which delayed the onset of reproduction.

Fecundity and adult lifespan had comparatively negligible effects on *r*_max_, which suggests that the carry over effect of reduced size at maturity on *r*_max_ is relatively weak. For example, at high-resource supply, adult lifespan and body size were greater at 26°C than at 32°C, yet fitness at 32°C was predicted to be 25% higher (Figs. 2c, 3). This is because high-resource supply and increased temperature minimised juvenile mortality and optimised development rate. This allowed faster recruitment at 32°C, leading to increased fitness as greater numbers of individuals could contribute to population growth through reproductive output sooner than for other treatments. This is consistent with general studies of ectotherm fitness [57], including mosquitoes [30]. This is a key finding, as it implies that predictions about the effect of warming on vector abundance and disease transmission based on laboratory-derived trait data (which are generally from populations under high-or optima resource supply) likely underestimate the effect of temperature on development time and juvenile survival, and overestimate effects of temperature on lifespan and fecundity.

Indeed, the trait-level responses of our high-resource supply treatments correspond with studies that have synthesised laboratory-derived trait responses to temperature to estimate vector fitness and *R*_0_. In these studies, the juvenile development rate of most mosquito vectors is expected to increase from ~0.07 day^-1^ at 22°C to ~0.14 day^-1^ at 32°C [8]. In the present study, development rate (1/development time, Fig. 1a) increased by a similar margin (~0.08 to ~0.17 day^-1^) across the same temperature range. In contrast, at low-resource supply, we found juvenile development rate was ~0.05 day^-1^ at 32°C (Fig. 1a). Such differences in juvenile trait responses are likely to substantially alter predictions about the temperature-dependence of *R*_0_. This underlines the importance of considering resource supply when predicting the temperature-dependence of *R*_0_ for vector-borne diseases.

Juvenile survival decreased significantly with temperature, and was overall significantly lower at low-resource supply (Fig. 1). This is probably because somatic maintenance costs increase with metabolic rate [21], which cannot be met below a threshold resource supply level. This explains why the highest level of mortality occurred at 32°C at low-resource supply, where the energy supply-demand deficit was expected to be the largest.

The Gompertz-shaped juvenile survival curves observed at 22 and 26°C at low-resource supply (Fig. 1a) may well arise from the amount of resource being sufficient for somatic maintenance, but not for development. This could be a key line of future investigation because it points to the importance of understanding how resource availability combines with temperature and other environmental factors to affect natural mosquito populations. For example, the negative effects of resource limitation on population growth through increased juvenile development time and mortality may be exacerbated, as individuals remain in the vulnerable juvenile stages for longer, which may increase predation threat [58]. If resource availability increases with climatic warming, the negative effects of predation on population growth could be offset by increased development and recruitment rates [59]. Alternatively, population growth could be dampened, if climate change reduces the quantity of food available to ectotherms [60,61].

We did not measure the effect of temperature and resource supply on fecundity directly, but used the size-scaling of this trait to estimate this effect. This is because most of the effect of resource limitation on juveniles is expected to affect adult mosquitoes indirectly by reducing size at emergence and lifespan [29,49]. Predicted fecundity increased nonlinearly with temperature, mediated by resource supply levels (Fig 1d, electronic supplementary material, Table S1). Across both resource levels, these fecundity estimates are similar to datasets that are used to parameterise mosquito-borne disease transmission models (e.g. [62]). However, even substantial under-or overestimation of fecundity by our size-scaling predictions and the use of starved adult lifespans, would not affect our main conclusions. This is because predicted fitness was relatively insensitive to these traits (Figs. 3, S3).

While the increased negative carry over effects of temperature at resource limitation on adult traits may have had a relatively weak impact on fitness compared to juvenile traits, temperature × resource supply interactions may have important effects on other components of vector-borne disease transmission [63]. For example, larger individuals may have greater transmission potential because they are more likely to outlive a pathogen’s external incubation period [64]. On the other hand smaller individuals may bite more frequently, which can increase transmission probability [65]. Also, larval nutrition [35] and temperature [66] can independently influence within-vector parasite development, but future studies could consider how the combined effects of temperature and resource supply affect this, and other important transmission traits.

In this study, we have not considered the temperature-dependence of resource supply itself (supply was held constant across temperatures in our experiments). In nature, the availability of resources may in fact be temperature-dependent. This is because microbial growth rates increase with temperature to some optimum, which may increase the concentration of food in the environment [9,67,68]. For example, *Anopheles* [69] and *Aedes* [70] mosquitoes can be reared exclusively on cultures of *Asaia* bacteria. We have also not directly addressed larval competition for resources by manipulating the number of larvae for a given resource supply level. Variation in larval density may introduce additional fitness constraints through interference and exploitative competition. It could also interact with temperature-dependent resource supply because a higher larval density will increase accumulation of waste products. These are interesting and potentially important avenues for future investigation.

We note that experimenting with more resource levels would not change our qualitative results, which we have shown to be robust using thorough sensitivity analyses. Indeed, the two resource levels we have chosen represent extremes, and it is reasonable to conclude that mosquito population fitness in the field fluctuates, with resource fluctuations, between the radically different temperature responses as we have found here (Fig. 3). One avenue for future work is to find more accurate methods to estimate effective temperature-dependent fitness values in the field, accounting for resource fluctuations.

Organisms experience significant resource limitation over space and time in nature. This is particularly true for insects such as mosquitoes, which have juvenile stages restricted to small, ephemeral aquatic habitats that are susceptible to resource fluctuations [17-19,71]. Our study underlines the importance of the effects of resource supply on the temperature-dependence of population-level fitness of an important disease vector. In doing so, our findings suggest that current projections of how climatic warming affects vector-borne disease transmission may prove inaccurate because they generally fail to consider resource limitations. Our findings also underline the need for future research effort to be directed at better understanding how temperature and resource supply interact in the field, and how this, and interactions between other environmental factors, may influence other components of vector-borne disease systems.

## Supporting information

Electronic Supplementary Information

