## Supplementary material for "The effect of resource limitation on the temperature-dependence of mosquito population fitness": Electronic Supplementary Information

### Electronic Supplementary Material

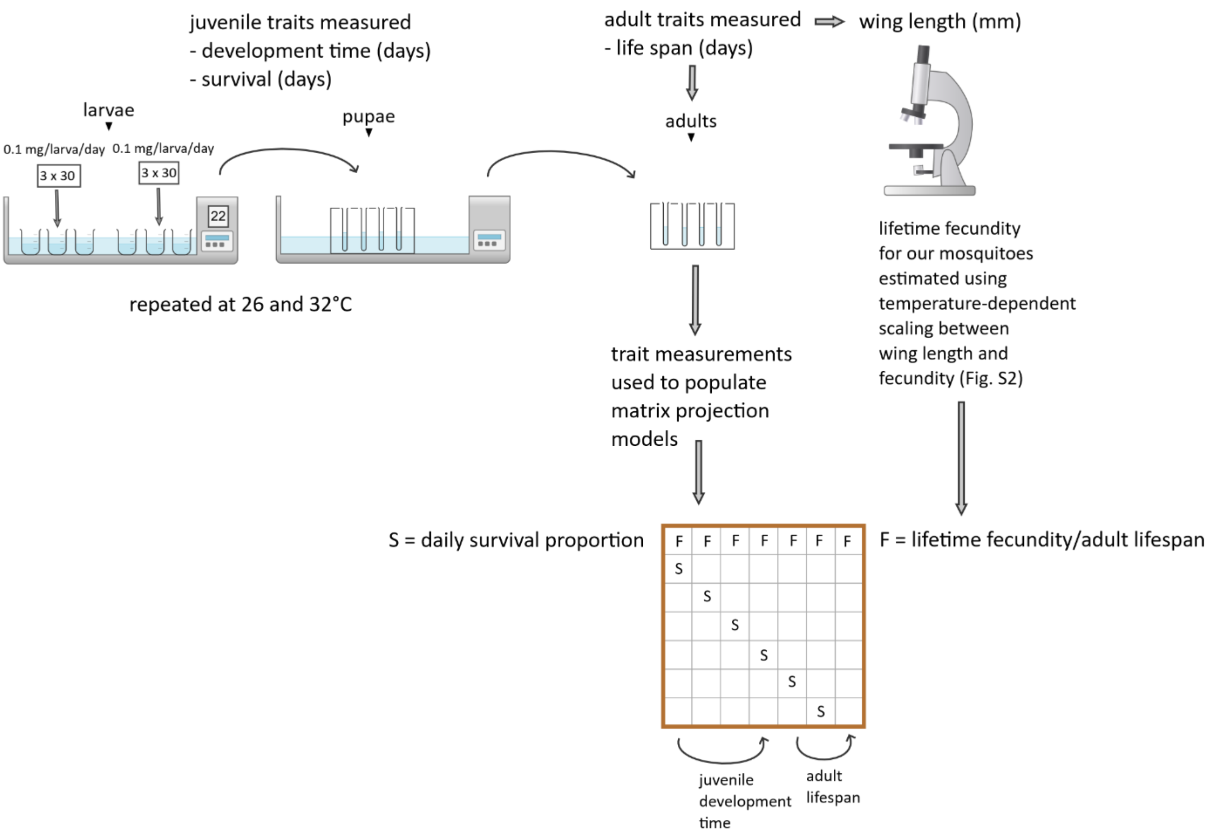


**Fig. S1** | A schematic of the experimental design and the traits measured. Juveniles were reared at three experimental temperatures (22, 26 and 32°C) and two resource supply levels (0.1 and 1 mg larva$^{-1}$ day$^{-1}$). We created three replicate tubs per treatment (90 larvae treatment$^{-1}$). Juvenile and adult traits were measured daily. Wing length was measured and used to predict individual lifetime fecundity (method described in the following section). Stage-structured matrix projection models were populated with these trait data to calculate $r_{max}$.

#### *Estimation of daily fecundity rate (first row of M in eqn. 1)*

We measured wing length as a proxy for body size and estimated fecundity using previously published datasets on the temperature- and resource supply-dependent scaling between wing length and fecundity [41,49]. Wing length (*L* in eqn. S1) was measured to the nearest 0.01 mm from the alula to the wing tip, excluding the apical fringe [72]. Wings (one per female) were removed, mounted onto glass slides, photographed using a dissecting microscope and then measured with ImageJ software [73].

Our analysis of the wing length-fecundity datasets [41,49] indicated that the effect of resource supply on fecundity occurs primarily through its effect on body size (electronic supplementary material, Fig. S2). So, to estimate lifetime fecundity (*F* in eqn. S1) from wing length for mosquitoes that we reared at 22 and 32°C, at high- and low-resource supply, we used the scaling coefficients from [41] at 20 and 30°C, respectively. For mosquitoes that we reared at 26°C, there is no corresponding temperature treatment in [41], so we used the scaling coefficients from another published study that was conducted at 27°C ([49], eqn. S1). Daily fecundity rate is required for the first row of **M** (eqn. 1), so lifetime fecundity was divided by lifespan and multiplied by 0.5 (assuming a 1:1 male-to-female offspring ratio) to give temperature-specific individual daily fecundity (*F* in Fig. S1).

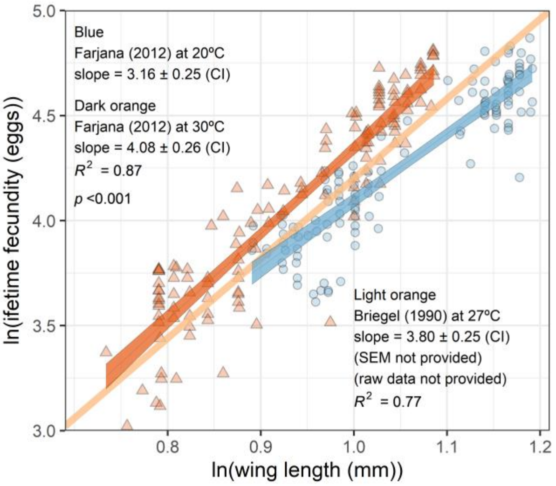


**Fig. S2 | a, Analysis of the Farjana et al.** [41] **dataset shows that the scaling of wing length and fecundity in *Ae. Aegypti* is temperature-dependent.** The scaling exponents (slopes) for both resource supply levels are significantly higher at 30°C than at 20°C. However, the effect of resource supply on fecundity is non-significant at the temperature level (not shown). The standard error for the scaling exponent at 27°C is not shown because it is not provided in [49], though the model's $R^2$ is satisfactory (0.77). Thus, to determine the sensitivity of $r_{max}$ to uncertainty in fecundity estimates at 26°C, we assumed a similar 95% CI (± 0.25) to those determined through our analysis of the Farjana et al. [41] dataset at 20 and 30°C (3.80 ± 0.25). Despite these assumptions relating to fecundity, our $r_{max}$ calculations are robust to uncertainty/variation in the underlying scaling and temperature dependences (Figs. 3 and S3).

$$\begin{aligned} &22°\text{C}, F = 0.93 + 3.16 \log(L) \\ &26°\text{C}, F = 0.40 + 3.80 \log(L) \\ &32°\text{C}, F = 0.26 + 4.08 \log(L) \end{aligned} \qquad \text{(S1)}$$

**Eqn. S1 | Scaling coefficients used to estimate temperature-dependent fecundity from wing length for our mosquitoes.** The coefficients come from our analysis of ([41,49], Fig. S2).

| Trait | Predictor | $\chi^2$ | df | *F* value | p-value |
|---|---|---|---|---|---|
| Juvenile development time | **Temperature** | **9.74** | **2** | **314.64** | **<0.001 *** ** |
| $R^2$ = 0.96 | **RS** | **50.30** | **1** | **3251.01** | **<0.001 *** ** |
| | **Temperature × RS** | **0.75** | **2** | **24.11** | **<0.001 *** ** |
| | Replicate | 0.01 | 2 | 0.18 | 0.83 |
| | Residuals | 2.30 | 148 | | |
| Lifespan | **Temperature** | **34.54** | **2** | **214.01** | **<0.001 *** ** |
| $R^2$ = 0.74 | **RS** | **3.18** | **1** | **39.42** | **<0.001 *** ** |
| | **Temperature × RS** | **2.41** | **2** | **14.95** | **<0.001 *** ** |
| | Replicate | 0.40 | 2 | 2.47 | 0.08 |
| | Residuals | 11.94 | 148 | | |
| Body size | **Temperature** | **0.65** | **2** | **111.58** | **<0.001 *** ** |
| $R^2$ = 0.79 | **RS** | **0.92** | **1** | **314.28** | **<0.001 *** ** |
| | **Temperature × RS** | **0.03** | **2** | **4.36** | **0.01 * ** |
| | Replicate | 0.003 | 2 | 0.48 | 0.62 |
| | Residuals | 0.41 | 141 | | |

**Table S1 | Type II Analysis of Variance results from GLMs fitted to the responses of life history traits to temperature and resource levels.** Significant effects are shown in boldface type. * ⇒ *p* value<0.05; ** ⇒ *p* value<0.01 *** ⇒ *p* value<0.001. Each pseudo $R^2$ (= residual deviance/null deviance) approximates how much variance the model was able to capture.

| Trait | Temperature (°C) | Mean ± s.e.m | |
|---|---|---|---|
| | | **Low-resource supply** | **High-resource supply** |
| | 22 | 36.45 ± 0.96 | 12.19 ± 0.25 |
| Development time (days) | 26 | 27.86 ± 0.64 | 8.53 ± 0.19 |
| | 32 | 21.00 ± 0.82 | 5.81 ± 0.14 |
| | 22 | 6.61 ± 0.39 | 11.35 ± 0.53 |
| Lifespan (days) | 26 | 5.59 ± 0.30 | 6.70 ± 0.35 |
| | 32 | 1.90 ± 0.17 | 2.46 ± 0.15 |
| | 22 | 2.65 ± 0.03 | 3.13 ± 0.03 |
| Wing length (mm) | 26 | 2.45 ± 0.02 | 2.93 ± 0.03 |
| | 32 | 2.28 ± 0.04 | 2.59 ± 0.03 |
| | 22 | 5.05 ± 0.83 | 4.29 ± 0.14 |
| Fecundity (eggs $female^{-1}$ $day^{-1}$) | 26 | 4.42 ± 0.25 | 6.79 ± 0.22 |
| | 32 | 12.25 ± 2.21 | 14.17 ± 1.47 |
| | 22 | –0.09 ± 0.02 | 0.17 ± 0.01 |
| Population-level fitness ($r_{max}$) | 26 | –0.03 ± 0.02 | 0.21 ± 0.03 |
| | 32 | –0.38 ± 0.06 | 0.28 ± 0.05 |

**Table S2 | Comparison of the effect of resource supply on the temperature-dependence of fitness and its component traits.** The means with standard errors for development time, lifespan and wing were estimated by using the GLMs in Table S1 (replicate dropped). For fecundity, these were estimated using the `Rmisc` package in R. For $r_{max}$, 95% CIs were approximated using the method described in [52].

| Model terms | Model name | AIC | ΔAIC | df |
|---|---|---|---|---|
| **Temperature × RS** | **Interaction** | **1050.85** | **0** | **7** |
| Temperature × RS + replicate | Full | 1053.05 | +2.2 | 9 |
| Temperature + RS | No interaction | 1053.50 | +2.7 | 5 |
| Temperature | Temperature only | 1055.99 | +5.1 | 4 |
| Resource | Resource only | 1110.30 | +59.5 | 3 |
| None | Null | 1108.70 | +57.9 | 2 |

**Table S3 | Simplification of the results of fitting the Gompertz juvenile survival model to the data**. The full fitted model includes the effects of temperature × resource supply and replicate on mortality. The interaction model without replicate won because it had the lowest AIC. ΔAICs were calculated as differences from the interaction model.

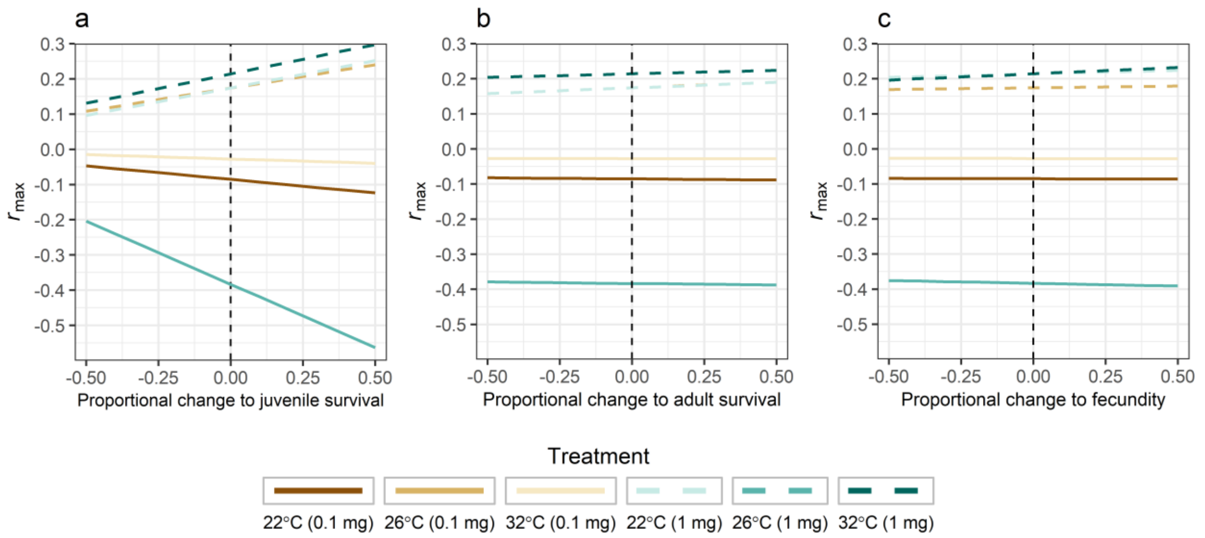


**Fig. S3 | Sensitivity of $r_{max}$ to proportional changes in juvenile survival (a) adult survival (b) and fecundity (c) for each *Ae. aegypti* population**. Juvenile survival was the most important contributor to $r_{max}$, as relatively small changes in the summed matrix elements for this trait would result in relatively large changes in $r_{max}$. Sensitivity of $r_{max}$ to adult traits was much weaker compared to sensitivity to juvenile traits. Dashed vertical lines denote $r_{max}$ at zero as represented by the midrange grey hue estimates in Fig. 3.

#### *Preliminary assays*

Prior to conducting the main experiment, we conducted a preliminary assay to determine the replication levels required to detect statistically significant effect sizes, and to calibrate our low-resource supply level (0.1 mg larva$^{-1}$ day$^{-1}$). Analysis of these data showed that our level of replication was sufficient; we detected a significant effect of temperature on juvenile development time and adult lifespan at low-resource supply at 26 and 32°C (Table S4). As an additional analysis, we compared the preliminary assay results with the corresponding treatment effects in the original manuscript. This analysis shows that there were no significant differences between these effects (Table S5).

**Table S4 | Type II Analysis of Variance results from GLMs fitted to the responses of juvenile development time and adult lifespan to temperature at resource limitation.** Significant effects are shown in boldface type. * ⇒ $p$ value<0.05; ** ⇒ $p$ value<0.01 *** ⇒ $p$ value<0.001. Each pseudo $R^2$ (= residual deviance/null deviance) approximates how much variance the model was able to capture.

| Trait | Predictor | $\chi^2$ | df | *F* value | p-value |
|---|---|---|---|---|---|
| Juvenile development time | **Temperature** | **0.73** | **1** | **26.36** | **<0.001 *****|
| $R^2$ = 0.48 | Replicate | 0.05 | 2 | 0.95 | 0.40 |
| | Residuals | 0.86 | 31 | | |
| Lifespan | **Temperature** | **5.54** | **1** | **113.43** | **<0.001 *****|
| $R^2$ = 0.76 | Replicate | 0.08 | 2 | 0.80 | 0.46 |
| | Residuals | 1.51 | 31 | | |

**Table S5 | Comparison of the pilot and the manuscript data on the effect of resource limitation on the temperature dependence of development time and adult lifespan.** Development time and lifespan were significantly lower at 32°C than at 26°C in both datasets. Treatment-level means with their standard errors for were estimated using the GLMs in Table S1 with replicate omitted. Overlapping confidence intervals (mean ± 1.96 × s.e.m) indicate that differences observed between the datasets for these traits were non-significant.

| **Trait** | **Temperature (°C)** | **Mean ± s.e.m.** | |
|---|---|---|---|
| | | **Low-resource supply** *Pilot* | **Low-resource supply** *Manuscript* |
| Development time (days) | 26 | 26.14 ± 1.04 | 27.86 ± 0.64 |
| | 32 | 19.38 ± 1.00 | 21.00 ± 0.82 |
| Lifespan (days) | 26 | 4.82 ± 0.30 | 5.59 ± 0.30 |
| | 32 | 1.99 ± 0.16 | 1.90 ± 0.17 |
